# ATP-induced asymmetric pre-protein folding: a driver of protein translocation?

**DOI:** 10.1101/202150

**Authors:** Robin A. Corey, William J. Allen, Ian Collinson

**Affiliations:** School of Biochemistry, University of Bristol, University Walk, Bristol, BS8 1TD

**Keywords:** SecA, SecYEG, protein translocation, molecular dynamics

## Abstract

The transport of proteins across membranes is a fundamental and essential process, achieved in every cell by the ‘Sec’ translocon. In prokaryotes, SecYEG associates with the motor protein SecA to carry out ATP-driven pre-protein secretion – a vital step in the biogenesis of most periplasmic, outer membrane and secreted proteins. Structural data of the SecA-SecYEG complex has provided considerable insight into underlying mechanism of this process. Previously, we have proposed a Brownian ratchet model for protein translocation, whereby the free energy of ATP binding and hydrolysis favours the progression of pre-protein across the membrane from the cytosol toward the outside [Allen, Corey *et al. eLife* 2016]. Here, we use atomistic molecular dynamics simulation of a SecA-SecYEG complex engaged with preprotein to further address the mechanism underlying this process. The data describe pre-protein secondary structure formation within the channel, which exhibits a nucleotide-dependent asymmetry between the cytoplasmic and exterior cavities. The results suggest ATP-dependent pre-protein transport is partly driven by pre-protein secondary structure formation. The model previously described, and refined here, could easily be adapted for the transport of proteins across various other membranes, such as the endoplasmic reticular and mitochondrial inner membranes.

## Introduction

The encapsulation and compartmentalisation of cells has necessitated the evolution of machineries which permit the transport of proteins across membranes, including for protein secretion and organellar import. Usually, protein transport occurs before the nascent protein has folded. This paper explores how the folding process *per se* may be exploited to drive protein translocation for the purposes of secretion.

The bulk of protein secretion in every cell is conducted by the ubiquitous Sec translocon, which also acts as the principle route for the insertion of membrane proteins. In bacteria, the translocon comprises SecY, SecE and usually SecG, wherein the protein-conducting pore runs through the centre of SecY. This complex can associate with either the ribosome for co-translational protein translocation (1) – the main pathway for nascent membrane protein insertion in bacteria (2, 3) – or with the motor protein SecA for post-translational secretion (4). In the latter case the fully synthesised protein is maintained in an unfolded conformation by chaperones, such as SecB, and SecA itself (4, 5). The protein must then fold during or after the translocation process.

Post-translational translocation of the unfolded pre-protein occurs through a contiguous channel formed through SecA and SecY (Figure 1A-C; (6-9)). The first step in this process is the ATP-dependent ‘initiation’ phase; whereby the pre-protein is targeted *via* its cleavable N-terminal signal sequence to SecA, and subsequently to the SecYEG complex. Transfer of the signal sequence to SecY unlocks the protein-channel enabling the intercalation of the mature protein (10, 11). Next, the pre-protein is fed through the SecY channel in a process driven by both ATP and the proton-motive-force (PMF) (12). Finally, the signal sequence is cleaved and the preprotein is either folded or trafficked onwards to the cell envelope or beyond that to the external medium.

**Figure 1:**
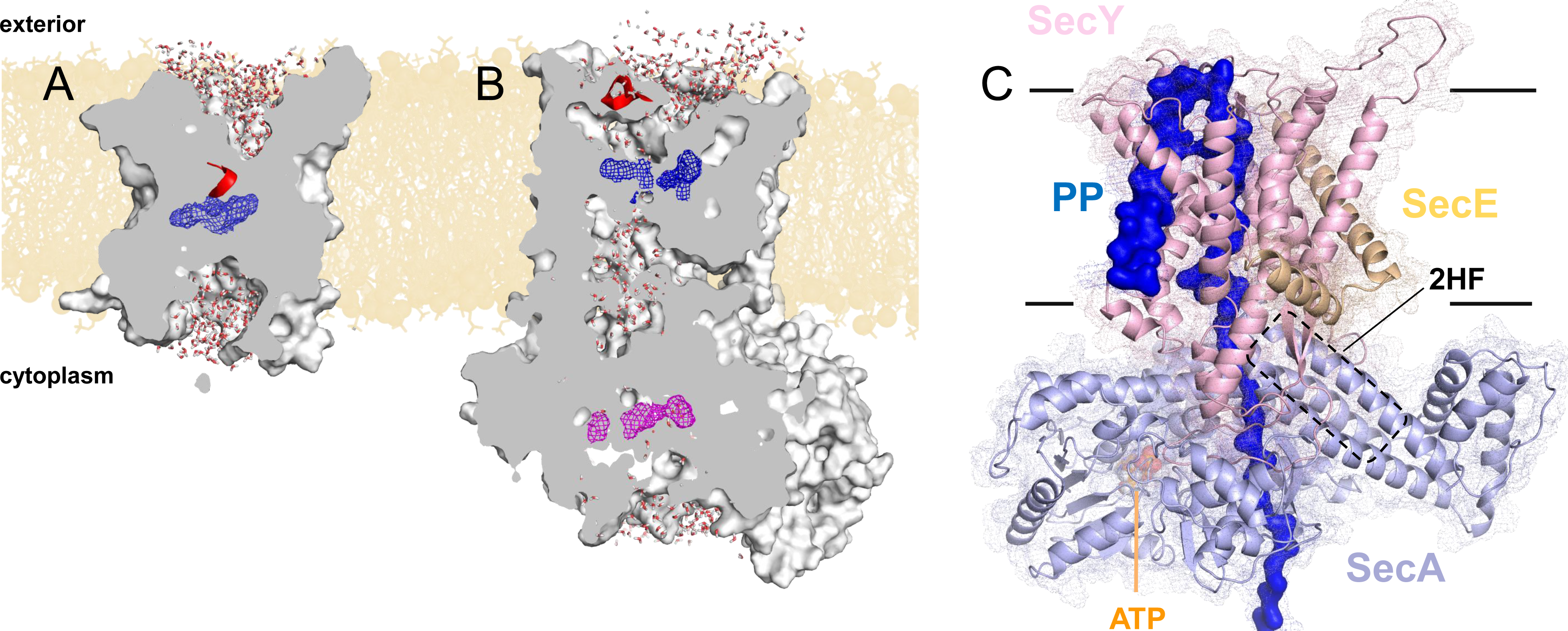
Structures of SecYEG and SecA. Views of A) SecYEβ (PDB 1RHZ; (7)) and B) SecYEG-SecA (PDB 3DIN; (6)) showing the cavities through the channel, with the protein in grey surface, the pre-protein pore constrictions in blue (SecY) or purple (SecA) mesh, and the SecY plug in red helix. The image was produced by embedding the crystal structures in a POPC membrane, solvating with explicit waters and allowing the non-heavy atoms to relax through restrained molecular dynamics (MD) over 4 ps. Degree of solvation and water density in the channel to be considered for illustrative purposes only. C) Cartoon of SecA-SecYEG with an engaged pre-protein, modelled from PDB 5EUL SecA (8). The Sec subunits are shown as cartoon with mesh overlay, with SecY in light pink, SecE orange, and SecA light-blue, with the 2HF highlighted. The unfolded pre-protein is shown in dark blue, labelled ‘PP’. The ATP analogue is coloured as orange, blue and red spheres. The approximate position of the membrane is marked.

The SecY pore is constricted centrally by a ring of six hydrophobic residues (7) (Figure 1A; blue mesh), which is flanked on the cytoplasmic and exterior sides by large solvated cavities, resulting in an hourglass-like shape (7) (Figure 1A). This ring is capped by the so-called ‘plug’ motif (7), which sits on the external face of the pore (Figure 1A; red helix) and helps to seal the channel against undesirable ion flux (13). Upon SecA binding, SecY rearranges to form a more open state (6), with this conformational switching likely controlled by the ATPase activity of SecA (14). This state still contains solvated cytoplasmic and exterior cavities, but the outwards movement of the plug (Figure 1B) and the insertion of the two-helix finger (2HF; Figure 1C) of SecA into the edge of the SecY cytoplasmic cavity profoundly changes the shape of the channel (Figure 1B). The 2HF has previously been shown to be coupled to the ATP binding region of SecA (the nucleotide binding site; NBS) (14-16), suggesting a nucleotide-dependent regulation of the SecY pore shape.

Recently, a pseudo-translocation state was captured by a crystal structure of *Geobacillus thermodentrificans* SecYE bound to *Bacillus subtilis* SecA (8). This was achieved by engineering a region of pre-protein into the SecA 2HF such that it resides within the SecY channel as a pre-protein would do. Amongst many observations, the structure reveals that the central pore of SecY tightly clasps the pre-protein (Figure 1 – figure supplement 1A).

We have previously proposed a Brownian ratchet mechanism for ATP-driven protein secretion, where stochastic pre-protein diffusion is biased into directional force by the action of the SecA ATPase (14). We demonstrated that this ATPase activity was able to open and close the SecY channel, thus allowing/ preventing passage of specific regions of pre-protein, such as large or charged residues, or regions of secondary structure. Feedback from the SecA 2HF permitted control of this mechanism, providing the means to bias Brownian motion (see Figure 8 and Video 1 from reference (14)).

Here, we present data extending this model to include structural changes within the channel and translocating pre-protein. Atomic models were built based on the pre-protein-containing crystal structure (8), remodelled into a physiological system. Molecular dynamics (MD) simulation data reveal key differences between the cytoplasmic and exterior cavities of SecY, with the cytoplasmic cavity widened appreciably when ATP is bound to SecA. This results in a substantially reduced degree of pre-protein secondary structure when compared to the exterior cavity. This asymmetry is not observed in the ADP-bound state, suggesting that pre-protein transport is, in part, driven by ATP-dependent control of secondary structure formation.

## Results

### Asymmetric secondary structure formation of translocon-engaged pre-protein

The pseudo-translocation crystal structure (PDB 5EUL (8)), described above, was remodelled into a physiological complex (*i.e.* a non-fusion protein) containing SecA, SecYE and 76 residues of an unfolded pre-protein, hereafter referred to as SecA-SecYE-PP; the details of this remodelling are described in Figure 1 – figure supplement 1B-C and Table 1. This structure was built into a solvated POPC bilayer, and simulated over 1 μs with the NBS of SecA occupied with either ADP or ATP. The simulations were stable within the core SecY region (Figure 2 – figure supplement 1) and, crucially, the SecY pore remained tightly formed around the bound pre-protein (Figure 2 – figure supplement 2A).

**Figure 2:**
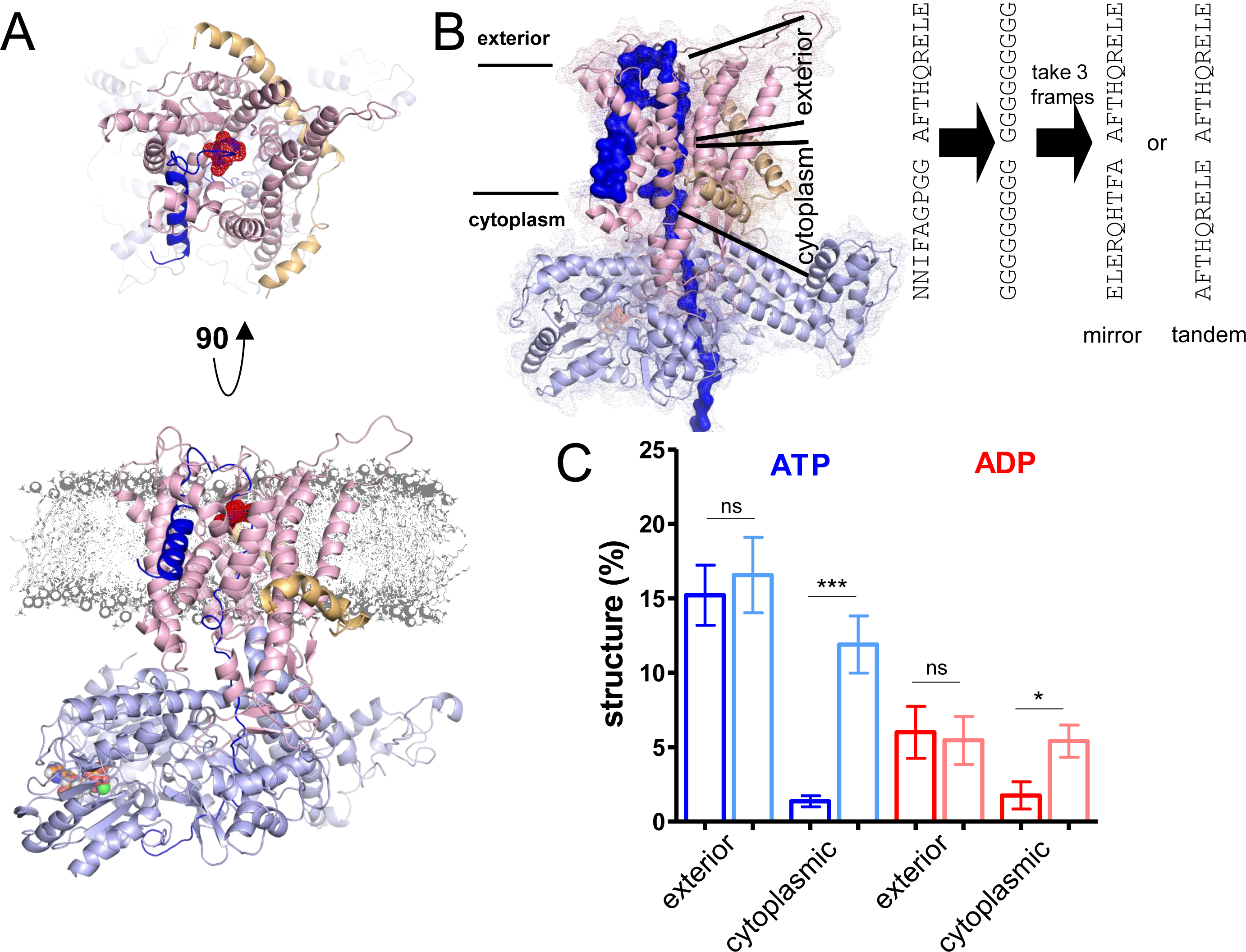
ATP-dependent asymmetric folding of pre-protein in the Sec translocon. A) Cartoon representation of a 1 μs MD snapshot of the SecA-SecYE-PP complex in the ATP-bound state, coloured as per Figure 1C. A region of α-helix, as computed by the dictionary of secondary structure of proteins (DSSP) and confirmed visually, is highlighted in red mesh. The approximate position of the membrane is shown in grey. B) Showing the work-flow to produce the pre-protein folding data. An 18-residue stretch of pre-protein is substituted to 18-Gly, then to a 9 residue stretch from the input model, repeated either mirrored or in tandem. The simulation times for each step are shown above. C) The degree of pre-protein folding in the exterior and cytoplasmic cavities of SecY in the ADP and ATP states. Shown are the combined datasets for the mirrored and tandem substrate (dark blue, red) as well as folding data for the same substrates in bulk water, i.e. not in the presence of translocon (light blue and pink). In both the ATP and ADP simulations the cytoplasmic cavity is significantly different from the bulk water in terms of pre-protein secondary structure (*p* < 0.0001 and *p* = 0.0135 respectively, from two tailed *t*-tests), although substantially more asymmetric in the ATP simulations. Reported *p* values are 0.675 and 0.822 for the exterior cavity. Error bars are s.e.m.

The most striking detail from the simulation data regards secondary structure within the pre-protein. In the region of the SecY channel, a range of hydrogen-bonded secondary structure motifs are observed, including considerable α-helix and 310-helix in the ATP and ADP simulations respectively (Figure 2 – figure supplement 2B). Visual inspection reveals considerable cross-pore asymmetry in this regard, with the majority of the secondary structure located in the exterior cavity (Figure 2A). Measuring the end-to-end distance of pre-protein in the channel – *i.e.* between the entrance to SecA and the tip of the exterior loop – reveals an average of ~0.18 nm per residue in both cavities for the ATP and ADP simulations. This value means that there is sufficient freedom for essentially all of the residues to adopt a helical configuration (based on ~0.15 nm and ~0.2 nm for residues in an α-helix or 3_10_-helix respectively), so secondary structure is not being disallowed due to restrictions in length in the input model.

### Quantifying cross-translocon pre-protein secondary structure

To better quantify the degree of pre-protein folding, further simulations were devised, aiming to maximise the information obtainable within current computational resource limits. Whilst much faster than tertiary structure folding, secondary structure formation is still reasonably slow with respect to atomistic simulation. Analysis of the initial trajectories (Figure 2 – figure supplement 2B) suggests that a time scale of ~100 ns is sufficient to kinetically sample helix formation. This matches time scales previously reported for model protein secondary structure formation (17); although it should be noted that any potential slower folding events, in the μs range, will not be sampled.

To provide broad conformational sampling and reduce any input bias from the starting configurations, multiple simulations were run from different starting coordinates. For this, new systems were modelled based on snapshots of the ADP- and ATP-bound systems at 1 μS. To abolish all of the secondary structure within the pre-protein, the 9 residues on each side of the central constriction were substituted with glycine and simulated for ~100 ns (Figure 2B), with the loss of structure confirmed using the dictionary of protein secondary structure (DSSP) algorithm (18, 19) (Figure 2 – figure supplement 3). Three snapshots that reported no secondary structure in this 18 residue region (Figure 2 – figure supplement 3 – black circles) were chosen from each simulation for further analyses. For each of these snapshots, the 9 residues on each side of the central constriction were substituted with the same sequence (AFTHQRELE) in a mirrored or tandem arrangement (Figure 2B). Then, 4-5 independent simulations were run for ~110 (+/-20) ns each, for a total of ~3 μS of simulation data of both the ATP- and ADP-bound systems.

Due to sampling restrictions we were realistically only able to analyse a single 9 residue sequence. We opted to use the sequence from the original crystal structure, as this has already been shown to be stable in the SecY channel, thereby reducing the potential for artefacts arising from changes of sequence. Additionally, the sequence has already been shown to fold within a reasonable time frame (Figure 2 – figure supplement 2B), and the ability to trap this sequence experimentally raises the possibility of direct comparison with experiment.

The simulations for each nucleotide state were analysed for hydrogen-bonded secondary structure in the 18 residue stretch of pre-protein through SecY. In the ATP-bound simulations there is >10 fold more pre-protein structure apparent in the exterior cavity than the cytoplasmic cavity (Figure 2C; blue data), despite the preprotein in both cavities being long enough to form considerable amounts of helix (Figure 2 – figure supplement 4A). Very little asymmetry with respect to pre-protein secondary structure is observed in the ADP-bound data (Figure 2C; red data). This suggests an ATP-dependent and highly asymmetric influence of the translocon on the folded state of the pre-protein; set up to promote folding on the exterior side of the channel.

### Pre-protein folding asymmetry is determined by the translocon

Simulations were conducted in order to ensure that the starting configurations of the pre-protein were not biasing the behaviour of the pre-protein (described in Figure 2B). The coordinates for the unfolded 18 residue stretch were extracted from the simulation and built into separate water boxes, without either SecA-SecYE or the membrane. Positional restraints were added to the ends of the peptide to replicate the restraints applied by the translocon and the bound signal sequence. These systems were then treated in the same way as the SecA-SecYE-PP simulations (simulated for 110 ns and each half analysed for secondary structure). The data show no significant asymmetry in pre-protein formation in either the simulation sets, be they derived from the ADP or ATP simulations (Figure 2C). Therefore, the asymmetric secondary structure formation must be a consequence of the translocon, rather than the intrinsic properties of the pre-protein. Comparison of the data with and without the translocon reveals no difference in the propensity for folding within the exterior cavity in either ADP or ATP simulations (Figure 2C). In the cytoplasmic cavity, a small difference is seen for the ADP simulations, compared to a very large difference in the ATP simulations (Figure 2C).

We next analysed the dependence of this effect on a correct ‘unlocking’ of the complex by signal sequence interaction (10). For this, we constructed a system in which the engaged pre-protein possessed a defective signal sequence (‘SS_Δ4_’; missing residues 5-9 (11, 20)), but retained its structurally-important position in contact with the lipid bilayer (11, 21, 22). Analysis of this complex as per the data in Figure 2C reveals that, whilst a cross-channel asymmetry is still observed in 2/3 of the simulation sets (Figure 2 – figure supplement 4B), the degree of pre-protein secondary structure is significantly higher in both cavities (Figure 2 – figure supplement 4C). Thus, a productive interaction of the signal sequence with the ATP-associated translocon is required to prevent pre-protein folding in the cytosolic cavity. In other words, the ability of the translocon to asymmetrically influence the folding propensity of the translocating pre-protein is subject to activation by ATP and the signal sequence.

### Perturbed translational and rotational dynamics of waters within the SecY translocon

The thermodynamics and kinetics of protein folding can be affected by perturbed water dynamics (23), and the waters within SecY have previously been shown to exhibit reduced mobility (24). To examine the role of water in the pre-protein folding process, 31 structural snapshots were extracted from each of the 1 μs SecA-SecYE-PP ATP and ADP simulations. Short simulations were run using a statistical ensemble which accurately models water dynamics (*i.e.* using an NVE ensemble – see Methods for details). For data analyses purposes only, a rectangular prism of 5 × 5 × 12 nm was imposed on each simulation box with its geometric centre on the SecY pore ring (Figure 3A). This prism was split into 24 slices of 0.5 nm along the z-axis, and the dynamics of the waters from each slice measured.

**Figure 3:**
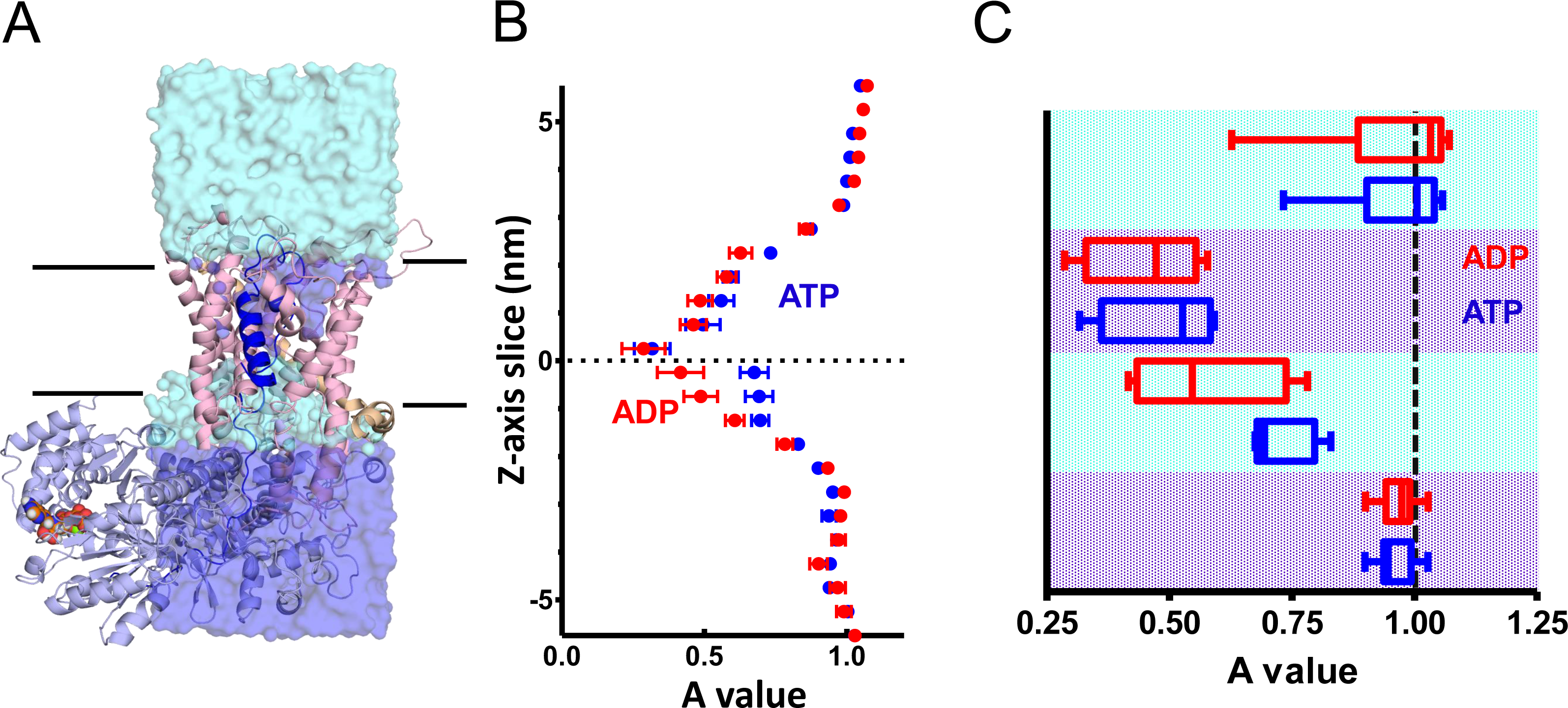
Perturbed water dynamics throughout the Sec translocon. A) Cartoon showing the prism of waters analysed for translational and rotational dynamics. The waters are shown as blue surface, alternating light and dark blue for the pooled data analysis in panel C. Note that whilst the membrane and solvent outside of this prism are missing from this figure, they were present in the simulation. B) MSD data of the waters along the length of the protein. The average MSD was calculated for each 0.5 nm slice, the data fitted to a power law equation, and the power value (A) for each slide was averaged across all 31 simulations. Here, the average for each slice is shown, with s.e.m as error bars. Both the ATP and ADP simulations are perturbed in the centre of SecY, but the ATP appears to be perturbed in an asymmetric manner. C) The values from panel B are pooled by region of the system, as illustrated by light and dark blue colouring in panel A, and the data plotted as box and whisker plots showing upper quartile, median and lower quartile.

To measure the translational water dynamics, mean squared displacement (MSD) calculations were employed (as applied in (24); see Methods for details). When applied to the waters through SecA-SecYE-PP, there is a clear pattern of perturbation in the centre of SecY, with the water dynamics at the centre of the channel severely restricted (Figure 3B and Figure 3 – figure supplement 1). Comparison of the waters on either side of the pore reveal an asymmetry in the ATP-bound system – *i.e.* a higher degree of translational diffusion in the cytoplasmic cavity than the exterior cavity – but not in the ADP-bound system (Figure 3C). Additional calculations were run for the translocon in a resting state (Figure 3 – figure supplement 2), which resembles the SecA-SecYE-PP in the ADP state, with no asymmetry between cavities. As with the pre-protein formation, these data mark the cytoplasmic cavity of the ATP-associated translocon as unusual in its environment.

In addition to the translational dynamics, rotational water dynamics were determined by computation of the corresponding dipole moments during the simulations. These data can be modelled using an autocorrelation function, C(t). As previously recorded (24), the waters at the centre of the translocon are too perturbed to accurately measure the rotational dynamics (Figure 3 – figure supplement 3).

### Folding of pre-protein and localised water dynamics in bulk solvent

To probe the apparent correlation between a decreased propensity for pre-protein secondary structure formation (Figure 2C) and increased translation water dynamics (and *vice versa*) (Figure 3B), we modelled peptide folding in constrained bulk solvent. For this, we applied positional restraints to the waters in a peptide-water simulation box, with the strength of the positional restraint chosen to simulate the reported MSD values for the water in the SecY cavities (cytoplasmic cavity = 21 kJ mol^−1^ nm^−2^; exterior cavity = 92 kJ mol^-1^ nm^-2^; Figure 3 – figure supplement 4A-B). Intriguingly, we see the opposite effect to that expected from Figure 2: the more ordered water results in a 4-5 fold decrease in secondary structure formation (Figure 3 – figure supplement 4C). This could plausibly be an effect of the higher-ordered water restricting the fluctuation (RMSF) of the peptide (Figure 3 – figure supplement 4D), *i.e.* the kinetics of pre-protein folding are slowed by ordered water, but not necessarily the thermodynamics. However, extension of 6 simulations for each condition to ~400-450 ns fails to change the observed effect (Figure 3 – figure supplement 4E).

These data suggest that ordering of water does not drive the folding process. An alternative explanation for the data is that the causation is reversed: pre-protein is causing the water ordering. To test this, a single folding trajectory (from Figure 3 – figure supplement 4C) that samples a range of unfolded and folded peptide configurations was taken. A number of snapshots along this trajectory were selected and equilibrated, with restraints on the protein but not on the water molecules. NVE simulations were run using the same restraints, and the computed water dynamics were compared to the recorded level of structure (Figure 3 – figure supplement 4F). Fitting a linear regression to the data reveals a statistically significant increase in water ordering with protein secondary structure, confirming that pre-protein folding is perturbing the water dynamics rather than the other way around. Thus, the perturbed water structure is likely to be partly a consequence of pre-protein folding, rather than a driving force for folding.

### Nucleotide-dependent variation in the geometry of the exterior and interior cavities of the translocon

The conformational entropy of a protein’s extended state is reduced when in a confined space, which should favour more compact, folded states (25). This raises the possibility that ATP-driven opening of the SecY translocon (14) might have a direct impact on pre-protein secondary structure. To test this, the cavity sizes of the translocons associated with either ADP or ADP were assessed throughout the 1 μs simulations. The pre-protein was removed from the translocon and the dimensions of the cavities on either side were measured for 31 structural snapshots from each simulation using the HOLE program (26). For consistency, the analyses were initiated at a set point between the residues Ile-78 and Ile-275 at the centre of the SecY pore (see Figure 1 – figure supplement 1A). In all of the simulation snapshots, a channel through SecY is found, and in many a channel through the whole complex can be traced (Figure 4 – figure supplement 1).

**Figure 4:**
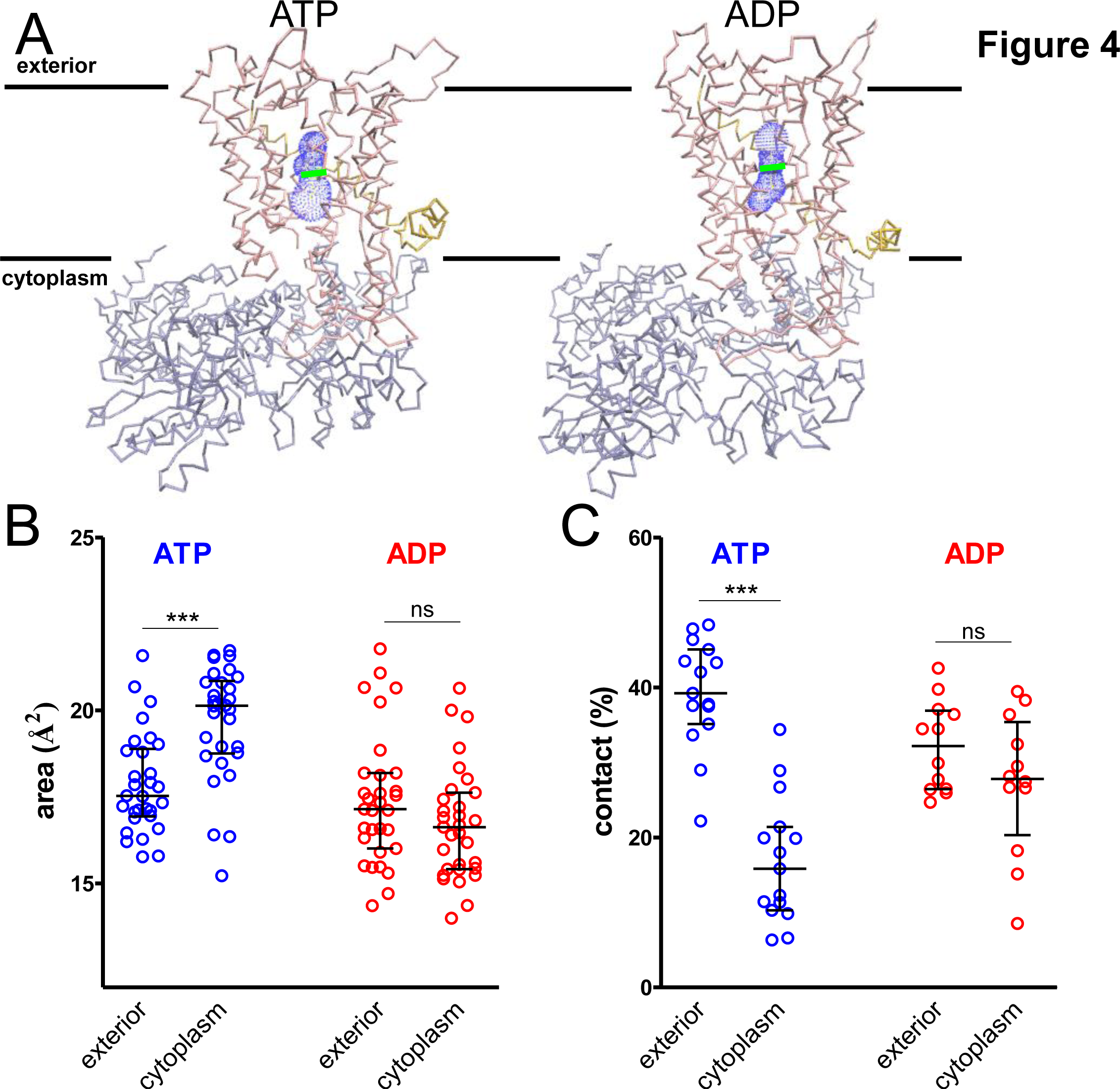
Perturbed water dynamics throughout the Sec translocon. A) Representative HOLE data used for averaging analysis. The protein backbone is shown as coloured trace and the calculated cavity as blue mesh. The central coordinate from the HOLE program is shown as a green line. On the left is the data at 608 ns of the ATP simulations, and on the right is at 610 ns of the ADP simulation. B) Quantified cavity sizes from the ATP and ADP simulation data. Plotted are the median and interquartile range. The difference between the exterior and cytoplasmic cavities was tested against an unpaired two-tailed *t*-test, reporting *p* values of <0.0001 and 0.1264 for ATP and ADP respectively. C) Degree of contact between the pre-protein and SecY channel, defined as interresidue distances of less than 0.3 nm, averaged over all of the residues. Correlated with a wider cavity, there is significantly less contact in the cytoplasmic cavity in the ATP-bound state (*p* <0.0001) but not in the ADP-bound stage (*p* = 0.1264).

The cavity volumes were measured from the constriction at the centre of the SecY pore and 6.5 Å either towards the cytosol (inside) or exterior (Figure 4A and Figure 4 – figure supplement 2A). The relative sizes of these defined regions reveal a strong asymmetry between cytoplasmic and exterior cavities in the ATP-bound simulations (Figure 4B; blue data) but not in the ADP-bound simulations (Figure 4B; red data). Once again, the ATP cytoplasmic cavity is the outlier, being at least 10% larger than any of the others. This in turn reduces the degree of pre-protein contact with SecY (Figure 4C), suggesting a role in SecY-pre-protein contact in mediating secondary structure formation.

Whilst it is difficult to deduce the precise mechanism for the ATP-induced increase in the cavity size of the cytoplasmic cavity, one candidate arising from the simulation data is the 2HF at the SecA-SecY interface (Figure 1C). The 2HF inserts into the SecY channel (6), contacts the translocating pre-protein (9, 27) and has a key, albeit disputed, role in protein translocation (14-16, 28, 29). In the simulation data, the distance from the tip of the 2HF to the central pore of SecY is substantially larger in the ATP state compared to the ADP state (Figure 4 – figure supplement 2B-C), providing a plausible mechanism for how the machinery regulates the size of its cytoplasmic cavity.

### Cavity size regulation occurs in the absence of pre-protein

To test whether the ATP-driven increase in size of the cytosolic chamber (Figure 4) is dependent on pre-protein, we analysed previously-produced simulation data (14) of the SecYEG-SecA complex (PDB code 3DIN (6)) with ATP and ADP bound, but without pre-protein. These systems proved challenging, as the HOLE program was unable to locate the correct cavity in 8/31 of the simulations of the translocon associated with ATP, and 20/31 of those associated with ADP. Comparison of the snapshots in which HOLE succeeded, however, revealed the same effect: *i.e* the cytoplasmic cavity of SecY is much larger than either the periplasmic cavity in the ATP-bound state or either cavity in the ADP-bound state (Figure 4 – figure supplement 3). Evidently then, the cytoplasmic cavity opens up in response to ATP irrespective of the presence of pre-protein.

### Secondary structure formation prevents transit through SecY pore

To determine whether the pre-protein secondary structures observed here impede passage through the SecY pore, a series of steered MD simulations were carried out. In these, a directional pulling force was applied to a stretch of the pre-protein through SecYE, with SecA and the rest of the pre-protein removed (Figure 5A). The pre-protein either contained an α-helical region – as sampled by equilibrium MD in the ATP-bound state (Figure 2), and stabilised here using a distance restraint between positions *i* and *i*+*4* – or had the helix abolished using a short steered MD simulation. These two substrates were pulled through the SecY pore, and the passage time of 10 independent repeats recorded. It was observed that the helix-containing substrate took considerably longer to pass (~10 times slower) than the substrate with no helix (Figure 5B). Although this setup clearly does not represent the physiological translocation process, the data indicate a significant barrier to the transit of folded regions of pre-protein through the protein channel.

**Figure 5:**
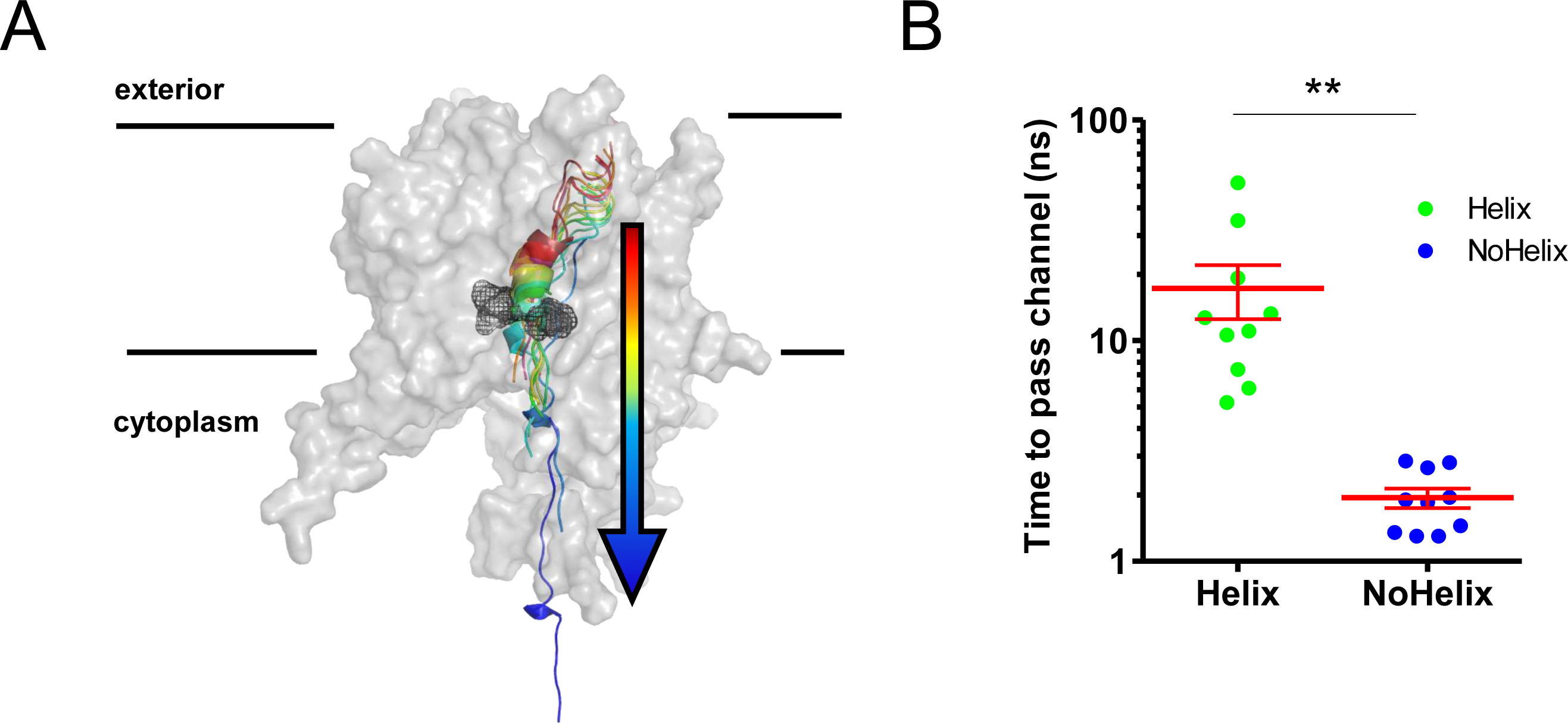
Pre-protein secondary structure prevents transit through SecY. A) Cartoon illustrating the principle behind the steered MD simulations. From a 1 μS snapshot of ATP-bound SecA-SecYE-PP, SecYE and a short region of helix-containing pre-protein were extracted, with the membrane trimmed to size. Two configurations of the pre-protein were established: either the helix was stabilised by a distance restraint between the hydrogen bonded atoms, or the helix was broken by a short steered MD simulation pulling the ends apart. A backward pulling force was then applied to each pre-protein, and the time taken to pass the channel measured B) Data from the simulations described in A. The presence of a helix in the pre-protein appears to be enough to restrict passage though the pore, *p* = 0.0045 using a two-tailed *t*-test.

**Figure 6: A model for pre-protein translocation through SecA-SecYEG**

In this model, SecA is shown in light blue, SecYEG in red, the SecY central cavity in orange and the pre-protein in dark blue. The direction of pre-protein diffusion is shown with a coloured arrow (green = forwards, red = backwards and orange = either).

The starting state (in the dashed box) represents the ADP-bound complex with no pre-protein secondary structure. In this state, the pre-protein is free to diffuse freely backwards and forwards, so no net translocation occurs. Then, localised pre-protein secondary structure formation can occur in either the exterior or cytoplasmic cavity; structure in the exterior cavity favours forward – productive – diffusion (top left panel), whereas structure in the cytoplasmic cavity would favour backward diffusion (bottom left panel). We previously demonstrated that bulky pre-protein in the cytoplasmic cavity triggers nucleotide exchange (14), resulting in the bottom right state, where ATP is bound and the cytoplasmic cavity has widened. As shown in Figure 2, this state strongly disfavours secondary structure in the cytoplasmic cavity, driving the complex towards the middle right panel. Here, either ATP is hydrolysed, to return to the original state, or secondary structure forms in the exterior cavity (as per the top right panel), resulting in a forward diffusion of pre-protein.

In this way, ATP is acting to prevent the backward diffusion of pre-protein in the complex, by shifting the equilibrium away from the bottom (unproductive) states (yellow arrow).

## Discussion

While it is widely accepted that pre-proteins need to be unfolded during transport (5, 30), the degree of unfoldedness is somewhat vague. The general assumption is that the tertiary structure forms after the translocation process, but what of the secondary structure? Does it form prior, during or after protein transport? This study seeks to address this problem in the context of the mechanism of protein transport.

A high degree of secondary structure has been reported for extended pre-proteins as they exit the ribosome (31, 32), and many proteins fold further once through to the periplasm (33). Furthermore, previous studies have intimated that pre-protein can form secondary structure within the translocon (27, 34). Here, we have performed a set of computational analyses on a model of a SecA-SecYE complex engaged with pre-protein (SecA-SecYE-PP), aimed at elucidating the importance of pre-protein folding for the process of protein translocation.

Our data reveal a propensity for the pre-protein to form secondary structure in both the cytoplasmic and exterior cavities of the channel. Strikingly, when ATP – and to a far lesser degree ADP – is bound to SecA, this propensity is highly asymmetric: secondary structure is strongly disfavoured in the cytoplasmic cavity (Figure 2). This cavity also shows markedly increased water dynamics (Figure 3), an increased cavity size (Figure 4B) and a decreased degree of contact between the translocon and pre-protein (Figure 4C). This observation suggests a specific role for ATP in regulating the structure of the translocating pre-protein: ATP binding causes the SecY cytoplasmic cavity to expand, reducing the amount of secondary structure.

Steered MD simulations confirm that regions of secondary structure are much less likely to pass the narrow SecY pore (Figure 5). Taken together, these effects could contribute significantly to any translocation model that incorporates pre-protein diffusion within the channel (14, 15). In our previously proposed ‘Brownian ratchet’ model (14), ATP binding is triggered by the presence of bulky regions – most likely secondary structure – at the cytoplasmic entrance to SecY. If ATP binding simultaneously unwinds pre-protein secondary structure in the cytoplasmic cavity of the channel, while not affecting folding in the exterior cavity, it would further bias the direction of diffusion; a model of how this might work is shown in Figure 6. Enticingly, these pre-protein conformational changes may be central to many other protein translocating machines, including chaperonins, organellar protein importers and the secretion systems of bacterial pathogens

It should be noted that we have largely ignored pre-protein sequence for this study, due to sampling restrictions within MD. The model protein from which our sequence was derived, pro-OmpA, eventually folds into a β-barrelled conformation, so the formation of α-helices could be considered surprising. However, the impact of nonlocal factors on the secondary structure of proteins is well-established (35), so an α-helix could easily be formed temporarily then broken in the absence of the final β-contacts.

Experimental validations of the findings reported here are difficult to achieve, as the ADP-bound complex is difficult to stabilise for structural analyses and the complexity of the translocation process makes it hard to disentangle the relevant effects from altering the pre-protein sequence and thereby its physico-chemical properties. Nevertheless, we feel the data presented here are compelling, and reveal considerable insight into the translocation process. The dependence of the key observations on nucleotide and the signal sequence is also heartening. Models for Sec-mediated pre-protein translocation, as well as other unrelated protein transport systems, should take into consideration a possible input from pre-protein secondary structure formation as a potential driver for transport.

## Methods

### Molecular dynamics simulations

All simulations were run using GROMACS 5.0.4 or 5.1.2 (36). Models for the simulations were built using chains A, Y and E of the crystal structure 5EUL (8) as starting coordinates, with missing loops added using Modeller (37), either modelling unstructured loops or modelled based on previous coordinates of SecA-SecYEG (PDB code 3DIN; (6)). Due to the non-physiological arrangement of the SecA 2HF and the pre-protein substrate of the input model, the 2HF was modelled based on 3DIN, and the substrate was extended in an unfolded conformation through the SecA, *via* known crosslinking sites (9). Substrate building was done with PyMOL (38). In the SecA NBS, the ADP-BeFx molecules were replaced with either ADP or ATP (39). Alternatively, simulations were built using the *M. jannaschii* SecYEβ crystal structure (7).

The protein and solvent atoms were described in the OPLS all-atom force field (40), with the simulations being run in an OPLS united-atom POPC membrane (41). The protein-membrane structures were built into simulation boxes with periodic boundary conditions in all dimensions and solvated with explicit SPC water and sodium and chloride ions to a neutral charge and concentration of 0.15 M. For the 1 μs SecA-SecYE-PP and *M. jannaschii* SecYEβ simulations, the systems were energy minimized using the steepest descents method over 2 × 5,000 steps, then equilibrated with positional restraints on heavy atoms for 1 ns in the NPT ensemble at 300 K with the Bussi-Donadio-Parrinello thermostat (42) and semi-isotropic Parrinello-Rahman pressure coupling (43, 44). Production simulations were run without positional restraints with 2 fs time steps over 500-1 μS on the UK HPC facility ARCHER, using time awarded by HECBioSim. Bond lengths were constrained using the LINCS method. Non-bonded interaction cut-offs were calculated using the Verlet method, with the neighbour search list updated every 20 steps. Long-range electrostatic interactions were calculated using the particle mesh Ewald method and a cut-off of 1.0 nm was applied for van der Waals and short range electrostatic interactions. All simulations reached a steady state as judged by their root-mean-squared-deviation from the starting structure (Figure 4 – figure supplement 1).

### Modelling a defective signal sequence

To investigate the effect of signal sequence interaction on the system, we modelled in a known defective signal sequence (11, 20), where a conserved four residue stretch is removed (here, KKTA**<u>IAIA</u>**VALAGFATVAS). We modelled this in PyMOL from a 1 μs snapshot of the ATP-bound SecA-SecYE-PP system. To reduce input bias, we allowed the N-terminal KK to remain in contact with the lipid phosphate groups and simply removed the four residues from the protein. We then repositioned the flanking residues slightly to allow bond formation, and relaxed the system using energy minimization. We then simulated the complex out to >400 ns for further analyses.

### Secondary structure formation analyses

Formation of secondary structure was determined both using visual analysis and quantified using the dictionary of secondary structure of proteins (DSSP) algorithm (18, 19). This was implemented using the Gromacs utility do_dssp, which provides an estimation of secondary structure for each reside based on dihedral angle, including ‘no structure’, ‘α-helix’, ‘3_10_-helix’ and ‘hydrogen-bonded turn’.

### Extending simulations to investigate secondary structure formation

To reduce the likelihood of sequence-dependent bias in the simulation data, a stretch of 9 residues on either side of the SecY pore was modelled as a mirror image repeat or in tandem. The sequence used (AFTHQRELE) was chosen as this was the sequence from the initial crystal coordinates. This means that the sequence is already known to sit happily within the translocon, and makes comparison to experimental data more straightforward. Initially, this 18 residue stretch was modelled as poly-glycine using SCWRL4 (45) in order to destroy the exiting secondary structure. Following steepest descents energy minimization and 1 ns equilibration, production runs were carried out until all of the secondary structure within this region had been abolished, as according to DSSP analysis (19); ~110 ns for the ATP-bound complex and ~80 ns for the ADP-bound complex (Figure 2 – figure supplement 3). For each simulation, three time points (70, 80 and 110 ns for ATP and 50, 65 and 80 ns for ADP, with only the 50 and 65 ns simulations run for the tandem sequence simulations) were chosen which displayed no structure at all according to DSSP analysis (Figure 2 – figure supplement 3A). This represents a reasonably broad configurational sampling, and prevents the analyses being too biased by a specific starting configuration.

For each time point, the 18 residue region was modelled to either ELERQHTFAAFTHQRELE or ELERQHTFAELERQHTFA using SCWRL4 (45), minimised using steepest descents, equilibrated for 1 ns and simulated for between 90 and 110 ns. The final 75 ns of each simulation run were then analysed for secondary structure. Simulations were run, in part, on EPCC’s Cirrus HPC Service.

### Water dynamics simulations

To model the water dynamics in and around the translocon, a series of simulations were run using structural snapshots of the 1 μs SecA-SecYE-PP ATP simulation from Figure 2 as starting points (500, 502, 504, 506, 508, 510, 600, 602, 604, 606, 608, 610, 700, 702, 704, 706, 708, 710, 800, 802, 804, 806, 808, 810, 900, 902, 904, 906, 908, 910, 1000 ns). Water dynamics simulations were run in the NVE ensemble, meaning a constant number of particles, volume and energy were maintained throughout. The advantage of this is to avoid introduction of artificial perturbations through use of a thermostat or barostat. Simulations were run with a 1 fs time step, writing coordinates every 5 fs. The Verlet cutoff scheme was used with a buffer size of 0.001 kJ mol^−1^ps^−1^ to achieve proper energy conservation, as monitored through following the energy of the simulations. Simulations were run on the University of Bristol’s High Performance Computer, BlueCrystal.

### Mean squared displacement calculations

To model the translational dynamics of the waters in the system, mean squared displacement (MSD) calculations were employed. MSD is a common statistical mechanics measure of translational motion. It follows the progression of an atom in relation to a reference position as a product of time, revealing the extent of its exploration of 3D space. Analyses were carried out as described previously (24). Briefly, a box of 5×5×12 nm was built around SecY, with the geometric centre at the pore ring. This box was subdivided into 24 slices of 0.5 nm, and the waters from each slice were analysed separately. The MSD of the waters in each slice was plotted, and the last 25 ps was fitted to a power law equations (Equation 1).

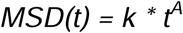

Where *k* is a fitting parameter, and the exponent *A* provides information on the molecular diffusivity. From this, the *A* value was plotted for each slice.

### SDAC calculations

To calculate the rotational dynamics of the waters, the same slices as used for the MSD calculations were subjected to SDAC calculations. Here, the data is modelled using an autocorrelation function C(t), which equals the degree of correlation of the dipoles at each time *t* in relation to the zero time point. For each data set, C(t) can be fitted to an exponential decay function, and the time at which the function reaches 1/e can be calculated, and plotted for each slice.

### Folding in bulk solvent with water position restraints

Waters dynamics in solvent boxes with 0.15 M NaCl were probed at a range of different water orders, as determined using MSD analyses. For this, restraining forces in the range of 0 to 1000 kJ mol^−1^ nm^−2^ were applied to the oxygen of each water molecule, and 100 ns simulations were run in the NPT ensemble, with isotropic pressure coupling. Snapshots were taken at 50, 60, 70, 80, 90 and 100 ns, and additional 50 ps simulations were run in the NVE ensemble. These simulations were analysed for translational water dynamics as described above, with the output plotted as a titration curve of restraining force vs. translational diffusion. The MSD values for the waters in the periplasmic and cytoplasmic cavities were read off of this curve.

Using the force constants representing water ordering in the periplasmic and cytoplasmic cavities, protein folding simulations were run of a peptide (ELERQHTFAAFTHQRELE) in an extended configuration. The setup of these followed the same technique as for the simulations in the main part of the study, except that equilibration was achieved through 10 ps NVT simulation with 0.5 fs time steps, 100 ps NVT simulation with 2 fs time steps and 100 ps NPT simulation with 2 fs time steps. Then, 100 ns production simulations were run, with secondary structure formation within the pre-protein analysed using DSSP.

### Folding in bulk solvent

Additional folding simulations were run and analysed as described above but without positional restraints on the water molecules, and instead of extended configuration for the peptide, coordinates were extracted from the simulations with the poly-glycine region in the translocon (Figure 3 – figure supplement 2). Position restraints were applied to the N and C termini of the peptide to reflect the restriction applied by SecA and the signal sequence, using 1000 kJ mol^−1^ nm^−2^ along the the z-axis and 10 kJ mol^−1^ nm^−2^ along the x and y-axes.

### HOLE

To analyse the SecY cavity sizes, snapshots were taken from the 1 μs SecA-SecYE-PP ATP and ADP simulations at 500, 502, 504, 506, 508, 510, 600, 602, 604, 606, 608, 610, 700, 702, 704, 706, 708, 710, 800, 802, 804, 806, 808, 810, 900, 902, 904, 906, 908, 910 and 1000 ns. In addition, snapshots were taken at the same time points from simulations previously published (14) of the SecA-SecYEG complex without pre-protein, with ATP or ADP bound.

For each snapshot, the centre of the SecY channel was initially defined as the geometric centre of the pore ring residues, and this was used to seed the HOLE calculations. The correct siting of the cavity was determined with visual inspection using VMD(46). For the successful calculations, data were extracted for 6.5 Å on either side of the pore ring. The area under the curves were computed and integrated using the trapezoidal rule with partitions of 0.5 Å.

### Steered MD

A 1 μs snapshot was taken from the SecA-SecYE-PP ATP-bound simulation, in which four pre-protein residues in the exterior SecY cavity form an α-helical configuration (Figure 2A). From this, SecA was removed and the backbone of the pre-protein was broken at residue Y1253 and removed from residue G1270 onwards (Figure 5A), and simulated for 5 ns. The α-helix was then either forced into an unstructured conformation using steered MD with a pulling force of 1000 kJ mol^−1^ nm^−1^ for 36 ps on residue E1260 or stabilised using a constraint of 0.2-0.25 nm between the backbone CO of residue 1262 and NH of 1266, followed by 5 ns unbiased simulation. Then, 10 independent steered MD simulations were run for each state, where the substrate was pulled from residue G1270 in a z-axis direction using a force constant of 600 kJ mol^−1^ nm^−1^. DSSP analysis confirmed that the helix remained formed or broken accordingly throughout the simulation. For each repeat, the time take for the helical/non-helical region to cross the pore was measured, based on the centre-of-mass distance between the region and the SecY pore residues, Ile-78, Ile-183, Ile-275 and Ile-404 (*G. thermodentrificans* numbering).

## Acknowledgements

This work was funded by the BBSRC: BB/M003604/1, BB/I008675/1 and BB/N015126/1 and Wellcome Trust: 104632. This work was carried out using the computational facilities of the Advanced Computing Research Centre, University of Bristol (http://www.bris.ac.uk/acrc/). Additional simulations were carried out using computer time on EPCC’s Cirrus HPC Service (https://www.epcc.ed.ac.uk/cirrus) and on the ARCHER UK National Supercomputing Service (http://www.archer.ac.uk), provided by HECBioSim, the UK High End Computing Consortium for Biomolecular Simulation (hecbiosim.ac.uk), supported by the EPSRC.

